# The effect of glucagon-like peptide-1 receptor agonist (GLP1RA) on hypertensive-induced heart failure with preserved ejection fraction and hypertensive cardiomyopathy

**DOI:** 10.1101/2023.02.12.528171

**Authors:** Zhe Yu Zhang, Song-Yan Liao, Zhe Zhen, Sijia Sun, Wing-Hon Lai, Anita Tsang, Jo Jo Siu-Han Hai

**Author notes:** Corresponding Author: Jo Jo Siu-Han Hai, MBBS (HK), MRCP (UK), Cardiology Division, Department of Medicine, The University of Hong Kong, Queen Mary Hospital, Hong Kong, China.

## Abstract

Emerging preclinical data suggest that glucagon-like peptide-1 receptor agonist (GLP1RA) possesses cardioprotective properties against the pathophysiology of hypertension (HT). We sought to unravel the potential therapeutic application of GLP1RA in a clinically relevant large animal model of hypertensive cardiomyopathy (hCMP). A combination of angiotensin II (Ang II) and deoxycorticosterone acetate (DOCA) pellets were used to induce sustained HT status and establish hCMP in porcine model. Changes in cardiac echocardiography, invasive hemodynamic parameters, neurohumoral biomarkers and inflammation-related cytokines were investigated in 23 adult pigs, among which 6 were serving as control, 9 were induced with HT, and the remaining 8 were HT-induced with GLP1RA treatment. Eight weeks after the study initiated, HT pigs have developed sustained high blood pressure (BP) at both systole and diastole. Phenotype of hCMP has also become significant as impairment in systolic/diastolic function, left ventricular remodeling and cardiac hypertrophy was determined by echocardiogram and invasive hemodynamics. Additionally, blood norepinephrine (NE) content, venoarterial NE gradient and pro-inflammatory cytokines in HT pigs were increased. GLP1RA treatment halted the elevation in BP, left ventricular remodeling and cardiac hypertrophy development; preserved the left ventricular systolic/diastolic function; reduced the venoarterial NE gradient as well as the pro-inflammatory cytokines at 18 weeks in pigs with hCMP. Our results demonstrate that GLP1RA treatment has a remarkable effect on BP decrease, inflammation suppression and left ventricular function improvement. Thus, we provide novel insight into the therapeutic potential of GLP1RA in HT-induced heart failure in a large animal model of hCMP.

## INTRODUCTION

Heart failure (HF) is a major health problem affecting over 23 million people worldwide (1). At least half of all patients admitted for HF are represented with HF with preserved ejection fraction (HFpEF) which is associated with significant morbidity and premature mortality (1). HFpEF differs from traditional HF with impaired systolic function or a reduced ejection fraction (HFrEF) by having a nearly normal systolic function with unaffected ejection fraction (EF) (2, 3). What features HFpEF is the impaired relaxation and increased diastolic stiffness which together give rise to ultimate ventricular diastolic dysfunction (4, 5). With unaffected EF, HFpEF is therefore associated with a better long-term survival and required a different therapeutic approach from HFrEF (6). However, better long-term survival leads to less research and clinical attention on HFpEF than HFrEF. Up-till-now, no existing pharmacological or device therapy has been shown to efficiently improve the clinical outcomes of HFpEF (7, 8).

Current existing difficulty in the development of effective therapeutic strategies for HFpEF is the insufficiency of a large animal model establishment in which the associated pathophysiology of HFpEF is simulated. Among various risk factors leading to HFpEF, chronic hypertension (HT) is highly associated and can develop in up to 80% or more of affected patients (9). Cardiomyopathy derived from HT or hypertensive cardiomyopathy (hCMP) is believed to largely contribute to diastolic dysfunction by impairing left ventricular (LV) diastolic relaxation and resulting in hypertensive LV morphological remodeling with cardiac hypertrophy under chronic high blood pressure (BP) burden (10, 11). Application of intravenous infusion of angiotensin II (Ang II) and subcutaneous implantation of deoxycorticosterone acetate (DOCA) mechanistically causes vasoconstriction and water-sodium retention which together increase the total peripheral resistance (TPR) and intravascular volume. According to known relationship between BP, TPR and cardiac output, both Ang II and DOCA are capable of elevating systematic BP (12). Previous studies have reported on regional heterogeneity in sympathetic nerve activity (SNA) in small animal model like rat induced with HT by administration of either Ang II or DOCA along with a high-salt diet (13, 14). Benefiting from the manageable size of small animal model, scientists have also discovered and established a strong connection between HT and the role of inflammation in HT-induced HFpEF (15, 16). Nonetheless, involvement of large animal model of HT with hCMP are limited over the past few decades (17–21), and the relationship between regional changes in SNA, systematic changes in inflammation and the development of HT remains elusive in large animal model.

In our study, we have successfully established a large animal model of sustained HT and hCMP in adult pigs that could mimic the clinical phenotype of refractory HT by using a combination of low-dose Ang II with a continuous infusion rate and low-dose DOCA with a constant release. Based on the porcine hCMP model created, we have investigated the potential therapeutic implications of glucagon-like peptide-1 receptor agonist (GLP1RA) in treatment of HT-induced HFpEF and hCMP.

## RESEARCH DESIGN AND METHODS

### Study Protocol

Our study course was divided by three timepoints: baseline, 8-week follow-up and 18-week follow-up. A total of 23 female farm pigs aging between 9-12 months and weighing between 35-45 kg were involved in our study. Accelerated HT with hCMP was successfully established in 17 animals using a combination of continuous Ang II infusion and subcutaneous DOCA pellets implantation. Remaining 6 animals were served as controls. A commercially available drug infusion pump (Tricumed Medizintechnik GmbH, Germany) which contained 120mg of Ang II in 12mL sterile isotonic saline was implanted subcutaneously into the ventral aspect of the animal’s right neck to induce HT in these animal models. The infusion pump was connected to an intravenous catheter inserted into the right internal jugular vein through a surgical cutdown. The rate that the infusion pump delivered Ang II was constant at 2.5uL/hr. The pump was refilled every four weeks to ensure a sustained delivery of Ang II. At the ventral aspect of the animal’s left neck, a subcutaneous pocket was surgically generated for DOCA pellets (Innovative Research of America, Sarasota, FL, USA) implantation. The release rate of DOCA from these pellets was 100mg/kg over 90 days. Extreme care should be taken to keep DOCA pellets away from any alcoholic solvents in order to generate the most optimal performance. DOCA pellets were designed to be absorbable, and refilling was not necessary throughout the entire experiment course. All 17 animals with developed HT at 8-week follow-up underwent selective angiogram of the celiac and superior mesenteric arteries, and 8 were selected to receive the treatment of a long-acting GLP1RA agent, Liraglutide, by randomization. Animals chosen to receive GLP1RA treatment were injected with a Liraglutide pen (Victoza, Novo Nordisk) subcutaneously around the neck region at a constant rate of 3mg/three-time-a-week.

A commercially available BP recording telemetered device (PhysioTel Digital telemetry D70, Data Sciences International, Saint Paul, MN, USA) was implanted in a pocket inside the neck region of all 23 pigs. This telemetered device was connected to an intra-arterial catheter which was inserted into the animal’s right carotid artery via a surgical cutdown to ensure a real-time BP measurement in conscious animals. After animals underwent general anesthesia (GA) with tiletamine and zolazepam (intramuscular injection of zoletil 20mg/kg), their echocardiography and invasive hemodynamics were investigated. In all animals, BP measurement was collected twice a week, echocardiographic assessments were done once-in-two-week, and invasive hemodynamic assessments were conducted three times at baseline, 8- and 18-week follow-up.

Local institutional ethics committee for animal research has approved our study protocol before experiment started. All experimental procedures were performed in strict accordance with the United States National Institutes of Health-published “Guide for the Care and Use of Laboratory Animals”.

Online-only Data Supplement provides detailed study methods of the invasive hemodynamic, echocardiographic, biomarker and histological assessments.

### Statistical Analysis

SPSS software (SPSS, Inc., Chicago, IL, USA) was used in the statistical analyses to allow us expressing the continuous variables as mean ± SEM and to perform different comparison tests. We used 2-way repeated ANOVA with Tukey test to compare the serial changes in BP, echocardiographic, invasive hemodynamic parameters and biomarkers at different time points between groups. Between 8- and 18-week follow-up, differences in BP, hemodynamic parameter and IL-6 were compared by 1-way ANOVA with Kruskal-Wallis test. Independent Student t test was applied to establish the comparison of immunochemical staining results between groups. Value of P<0.05 marked the significance in these statistical analyses.

### Data and Resource Availability

The data generated, analyzed and supported the findings of this study are available from the corresponding author on reasonable request.

## RESULTS

### Pressure Data

No significant changes were found in C group under the parameters of systolic, diastolic BP, LVESP and LVEDP from baseline to 8- and 18-week follow-up (**Figure 1A through 1D**; P>0.05). Upon action of the Ang II infusion and DOCA pellets implantation, systolic and diastolic BP successfully elevated in HT group and Tx group compared with C group at 8-week follow-up (**Figure 1A and 1B**; P<0.05). Interestingly, while systolic and diastolic BP kept increasing in HT group, these two indices showed a significant decrease in Tx group compared with HT group at 18-week follow-up (**Figure 1A and 1B**; P<0.05). Similar increasing trend of LVESP and LVEDP exhibited statistical significance in HT group and Tx group at 8-week follow-up and HT group alone at 18-week follow-up compared with its baseline and C group (**Figure 1C and 1D**; P<0.05). However, these pressure indices were decreased again in Tx group at 18-week follow-up, and the significant changes compared with its baseline or C group was absent (**Figure 1C and 1D**; P>0.05). Due to the reduction in both BP and LV pressure in Tx group, a significant decrease between HT group and Tx group at 18-week follow-up was observed (**Figure 1C and 1D**; P<0.05). Increase in systolic and diastolic BP in HT group indicated that we have successfully established a HT animal model, and the continuous elevation ensured a sustained HT by combination of Ang II infusion and DOCA pellets implantation. Increased pressure was also observed in terms of LV systole and diastole, nevertheless, all these pressure burdens loaded by HT were significantly relieved with GLP1RA treatment.

**Figure 1.**
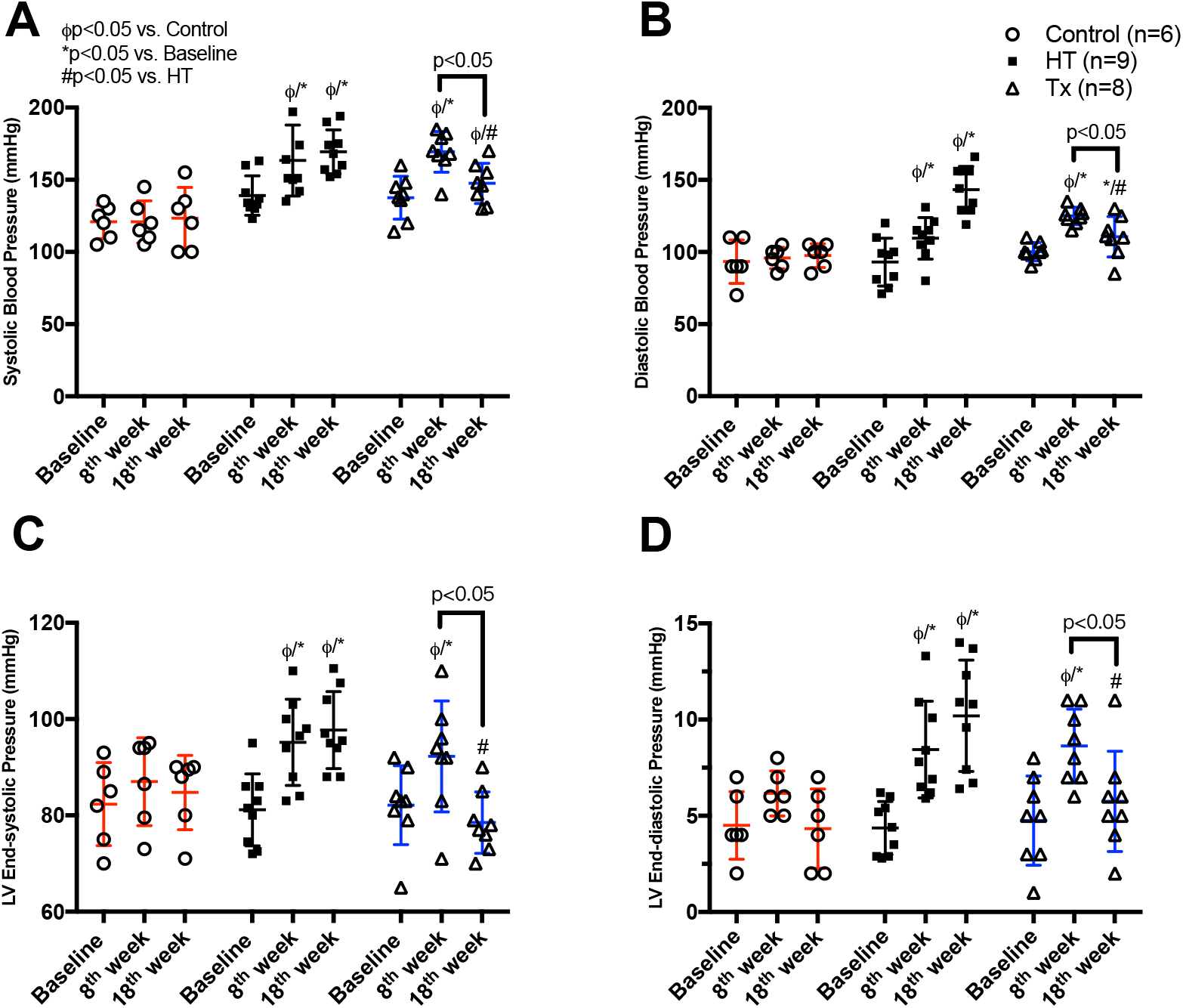
Hemodynamic parameters at baseline, 8-week and 18-week follow-up in control, hypertension and GLP1RA treatment groups. Serial changes to ambulatory blood pressure (**A and B**), left ventricular end-systolic pressure, and left ventricular end-diastolic pressure (**C and D**) in control (n=6), hypertension (n=9) and GLP1RA treatment (n=8) groups. ϕ<0.05 versus Control; *<0.05 versus Baseline; #<0.05 versus Hypertension.

### Invasive Hemodynamic Data

Detailed invasive hemodynamic assessments were performed using LV pressure-volume loop analysis (Figure S?). LV systolic contractile function as determined by +dp/dtMAX, end-systolic pressure volume relationship (ESPVR), -dp/dtMAX and end-diastolic pressure volume relationship (EDPVR) were similar in C group at 8- and 18-week follow-up compared with its baseline (**Figure 2A through 2D**; P>0.05). At 18-week follow-up, there was a significant decrease in +dp/dtMAX and ESPVR compared with 8-week follow-up in HT group (**Figure 2A and 2B**; P<0.05), suggesting a significant LV systolic function impairment. Nonetheless, this dysfunction was not shown in Tx group along the study course (**Figure 2A and 2B**; P>0.05). There was no significant difference in the index of -dp/dtMAX in either HT group or Tx group at both 8- and 18-week follow-up (**Figure 2C**; P>0.05), however, a LV diastolic function impairment shown by an increase in EDPVR was observed in HT group at 8- and 18-week follow-up compared with its baseline and C group (**Figure 2D**; P<0.05). In contrast, Tx group showed a significant decrease in EDPVR at 18-week follow-up from 8-week follow-up compared with HT group (**Figure 2D**; P<0.05). Gathering the results obtained from invasive hemodynamics, we found that developing HT is the key driving force in diastolic dysfunction yet GLP1RA treatment may provide therapeutic solution to HT-induced diastolic impairment.

**Figure 2.**
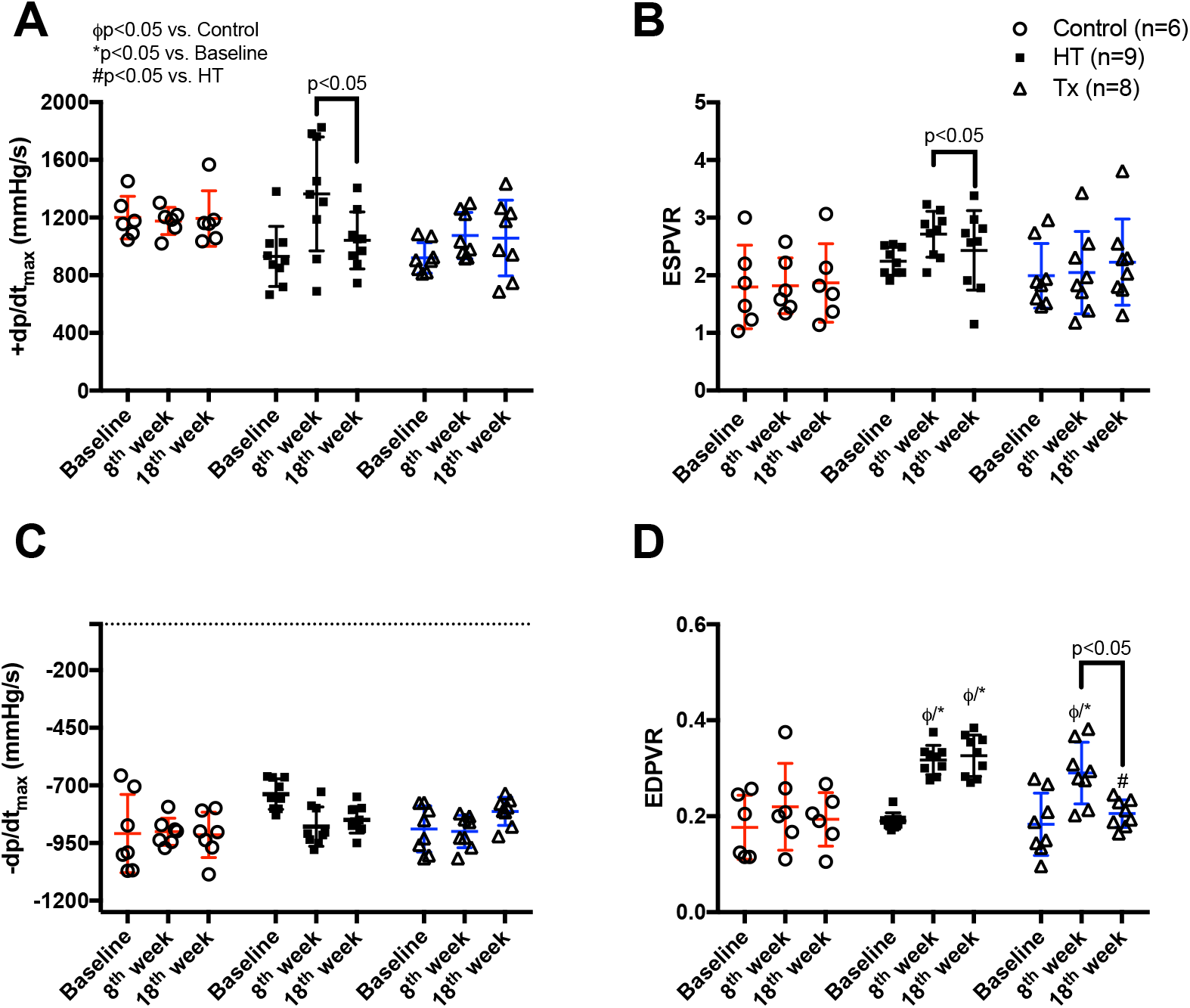
Pressure-volume loop assessment at baseline, 8-week and 18-week follow-up in control, hypertension and GLP1RA treatment groups. Serial changes to left ventricular systolic and diastolic function as determined by left ventricular +dp/dt (**A**), end-systolic pressure-volume relationship (**B**), left ventricular -dp/dt (**C**) and end-diastolic pressure-volume relationship (**D**) in control (n=6), hypertension (n=9) and GLP1RA treatment (n=8) groups. ϕ<0.05 versus Control; *<0.05 versus Baseline; #<0.05 versus Hypertension.

### Echocardiographic Data

In pigs from C group, there were no serial changes in echocardiographic measurements of IVSd, LVPW thickness, LV mass index, LVd at both systole and diastole, or LVEF from baseline to 8- and 18-week follow-up (**Figure 3A through 3E**; P>0.05). At 8-week follow-up in HT group and Tx group as well as 18-week follow-up in HT group alone, IVSd, LVPW thickness and LV mass index were significantly increased compared with their baseline and C group (**Figure 3A through 3C**; P<0.05). Differing from above, no significant changes were observed in LVPW thickness or LV mass index in Tx group at 18-week follow-up compared with C group (**Figure 3B and 3C**; P>0.05). Absence of significant changes were observed in LVd from all three group along the study course (**Figure 3D**; P>0.05). A decrease in the LVEF was detected in HT group when comparing itself at 8-week follow-up versus 18-week follow-up (**Figure 3E**; P<0.05), yet this significant reduction was not observed in C group and Tx group (**Figure 3E**; P>0.05), indicating that our animal models mimicked the phenotype of clinical HFpEF to a great extent. We additionally investigated LA volume and found no significant change in C group (**Figure 3F**; P>0.05), while HT group and Tx group at 8-week follow-up as well as HT group alone at 18-week follow-up exhibited statistically significant increase compared with its baseline and C group (**Figure 3F**; P<0.05). Results from echocardiographic assessment indicated that not only LV but LA has also been severely affected by sustained HT, and GLP1RA has the potential to attenuate the worsening in cardiac function to the contrary.

**Figure 3.**
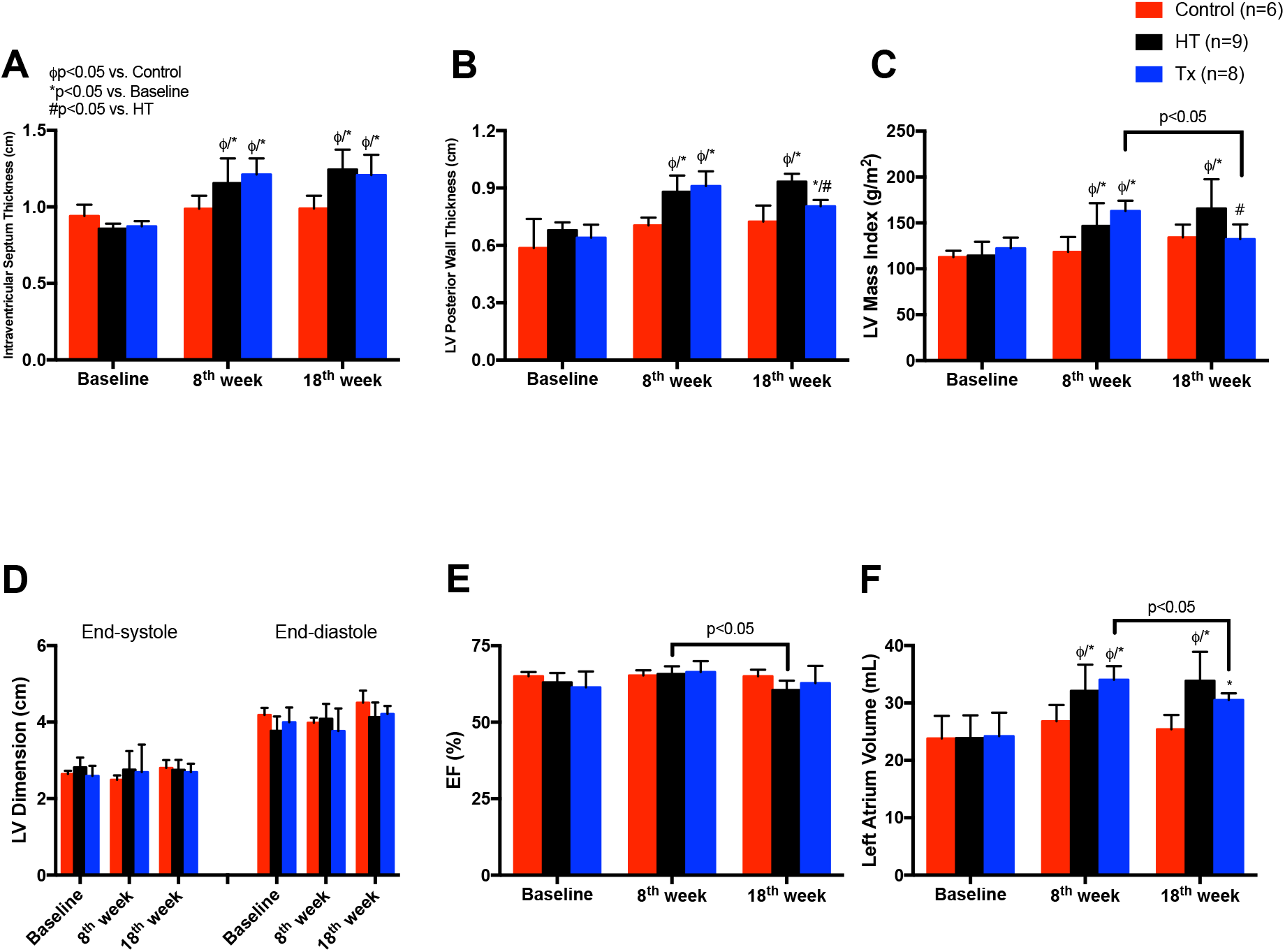
Echocardiographic parameters at baseline, 8-week and 18-week follow-up in control, hypertension and GLP1RA treatment groups. Serial changes to echocardiographic assessments of the left ventricle as determined by intraventricular septum thickness (**A**), left ventricular posterior wall thickness (**B**), left ventricular mass index (**C**), left ventricular dimension (**D**) and left ventricular ejection fraction (**E**), and serial changes to echocardiographic measurement of the left atrium as determined by left atrium volume (**F**) in control (n=6), hypertension (n=9) and GLP1RA treatment (n=8) groups. ϕ<0.05 versus Control; *<0.05 versus Baseline; #<0.05 versus Hypertension.

### General Body Condition and Anatomical Data

Since the animal models involved in our study were already adult pig model, body length in three groups did not show significant difference along the experiment course (**Figure 4A**; P>0.05). However, body weight between groups varied even though all pigs were fed with same chow diet and averagely same amount. HT group exhibited a continuous increase in body weight from baseline to 18-week follow-up, and the body weight measured at 18-week follow-up was significantly higher than its baseline and C group (**Figure 4B**; P<0.05). In Tx group, body weight at 18-week follow-up showed no significant difference compared with its baseline and C group (**Figure 4B**; P>0.05), yet a dramatic decrease was observed compared with HT group (**Figure 4B**; P<0.05). Given by body length data, the body weight-to-body length ratio (a make-up parameter mimicking body mass index in human) was significantly elevated in HT group at 18-week follow-up compared with its baseline, C group and Tx group (**Figure 4C**; P<0.05). Heart rate data were also collected in our study. Both HT group and Tx group revealed significantly increased heart rate at 8- and 18-week follow-up compared with their baseline and C group (**Figure 4D**; P<0.05).

**Figure 4.**
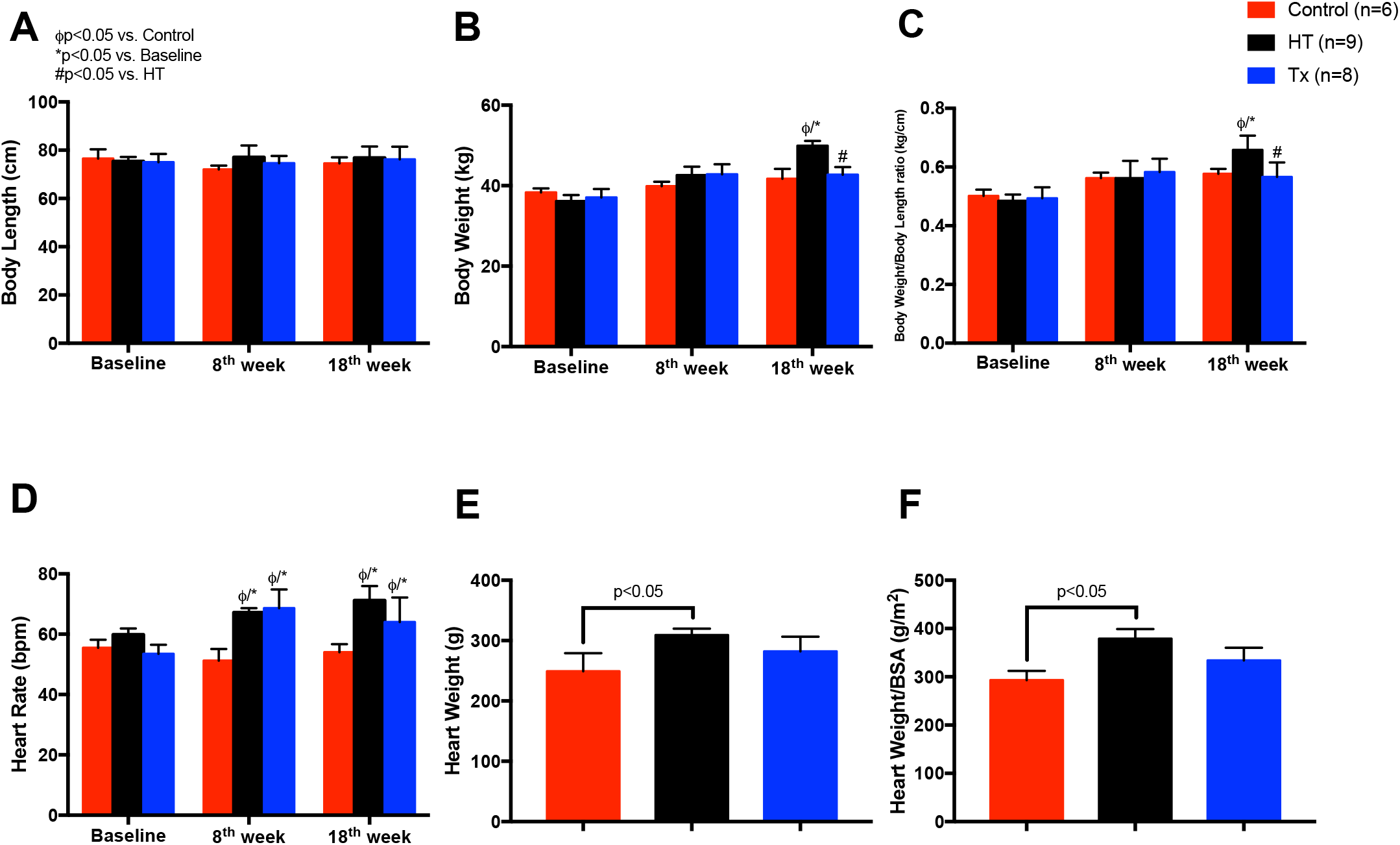
General body and heart condition, and heart rate at baseline, 8-week and 18-week follow-up in control, hypertension and GLP1RA treatment groups. Serial changes to body length (**A**), body weight (**B**), body weight/body length ratio (mimicking human body mass index; **C**) and heart rate (**D**) as well as heart weight (**E**) and heart weight/body surface area (**F**) in control (n=6), hypertension (n=9) and GLP1RA treatment (n=8) groups. ϕ<0.05 versus Control; *<0.05 versus Baseline; #<0.05 versus Hypertension.

Gross anatomy of LV (**Figure S1**) showed a significant increase in heart weight at 18-week follow-up in HT group compared with C group (**Figure 4E**; P<0.05), yet such increase was not observed between C group and Tx group (**Figure 4E**; P>0.05). Heart weight-body surface area ratio exhibited the same increase between HT group and C group (**Figure 4F**; P<0.05). General body condition and the heart anatomical results demonstrated that sustained HT has clearly caused body weight and heart weight increase, giving rise to hypertensive cardiac hypertrophy. Intriguingly, GLP1RA treatment in animals with developed HT might have the potential to attenuate the abnormal body weight gain and reverse the hypertrophy in LV. Of note, GLP1RA did not show with capability in decreasing HT-induced heart rate elevation to baseline as expected, and this is possibly due to the direct activation of GLP1RA receptor located on the sino-atrial node so that the heart rate stimulation follows (22).

### Histological and SNA Data

Masson’s trichrome staining (**Figure S2**) revealed a dramatic increase in percentage area of fibrosis at the LV endocardium and mid-myocardium excised from animals sacrificed at 18-week follow-up in HT group compared with C group and Tx group (**Figure 5A and 5B**; P<0.05). However, no significant fibrotic increase was observed at LV epicardium among three groups (**Figure 5C**; P>0.05). As expected, Ang II and DOCA induced hypertensive cardiomyopathy exhibited extensive fibrosis in LV, suggesting the development of HF with reduced numbers of functional cardiomyocytes.

**Figure 5.**
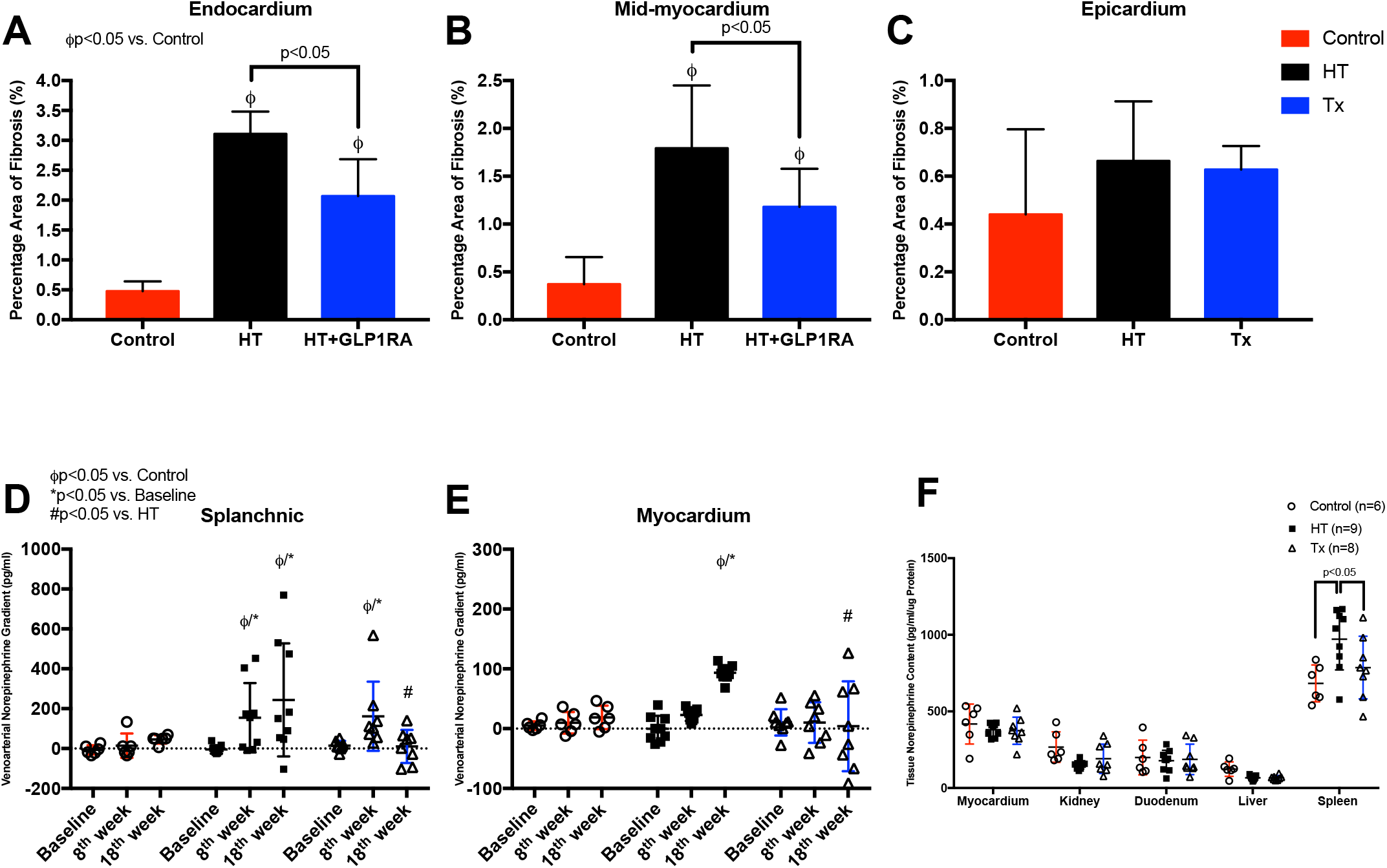
Myocardial fibrosis, venoarterial norepinephrine gradient and tissue norepinephrine content in control, hypertension and GLP1RA treatment groups. Percentage area of fibrosis in endocardium (**A**), mid-myocardium (**B**) and epicardium (**C**) as well as serial changes to venoarterial norepinephrine gradient over splanchnic organs and myocardium (**D and E**), and tissue norepinephrine content in various organs (**F**) in control (n=6), hypertension (n=9) and GLP1RA treatment (n=8) groups. ϕ<0.05 versus Control; *<0.05 versus Baseline; #<0.05 versus Hypertension.

In HT group, both splanchnic and cardiac venoarterial NE gradient got significantly elevated at 18-week follow-up compared with its baseline and C group (**Figure 5D and 5E**; P<0.05). The increase in splanchnic gradient already became statistically significant at 8-week follow-up (**Figure 5D**; P<0.05) while the cardiac gradient at 8-week follow-up did not show any significance (**Figure 5E**; P>0.05). At 8-week follow-up, Tx group exhibited significantly higher splanchnic gradient compared with its baseline and C group (**Figure 5D**; P<0.05). However, when splanchnic gradient comparison established between Tx group and HT group at 18-week follow-up, Tx group showed a statistical decrease (**Figure 5D**; P<0.05). No obvious changes were observed in the cardiac gradient of Tx group at wither 8-week follow-up or 18-week follow-up compared with its baseline and C group (**Figure 5E**; P>0.05); this gradient in HT group significantly elevated compared with Tx group at 18-w eek follow-up (**Figure 5E**; P<0.05). In terms of tissue NE content measured from myocardium, kidney, duodenum, liver and spleen, spleen was the only organ detected to have a significantly higher tissue content in HT group compared with C group and Tx group (**Figure 5F**; P<0.05). Since NE is one of the most important neurotransmitters in SNS, its hormone level expressed in either venoarterial gradient or tissue can reflect the systemic and local SNA, respectively (23). In sustained HT status, animals exhibited a continuous increase in venoarterial NE gradient as well as splenic tissue NE. Interestingly, GLP1RA treatment effectively repressed NE elevation, suggesting a SNA tune-down.

### Biomarkers and Pro-Inflammatory Cytokines Data

At 18-week follow-up, the plasma Ang II, plasma NE, plasma BNP and plasma adrenaline in HT group was significantly elevated compared with its baseline and C group (**Figure 6A and 6C**, **Figure S3A and S3B**; P<0.05). In addition, the plasma Ang II was also significantly increased in HT group compared with Tx group (**Figure 6A**; P<0.05). In Tx group, the plasma Ang II at both 8- and 18-week follow-up, the plasma NE and plasma adrenaline at 18-week follow-up exhibited significantly higher than its baseline and C group (**Figure 6A and 6C, Figure S3B**; P<0.05). Tx group also showed a significant increase in plasma BNP inter-group between itself at 8- and 18-week follow-up (**Figure S3A**; P<0.05). There were no serial changes detected in the plasma renin or plasma creatinine between all three groups at each timepoint (**Figure 6B and S3C**; P>0.05).

**Figure 6.**
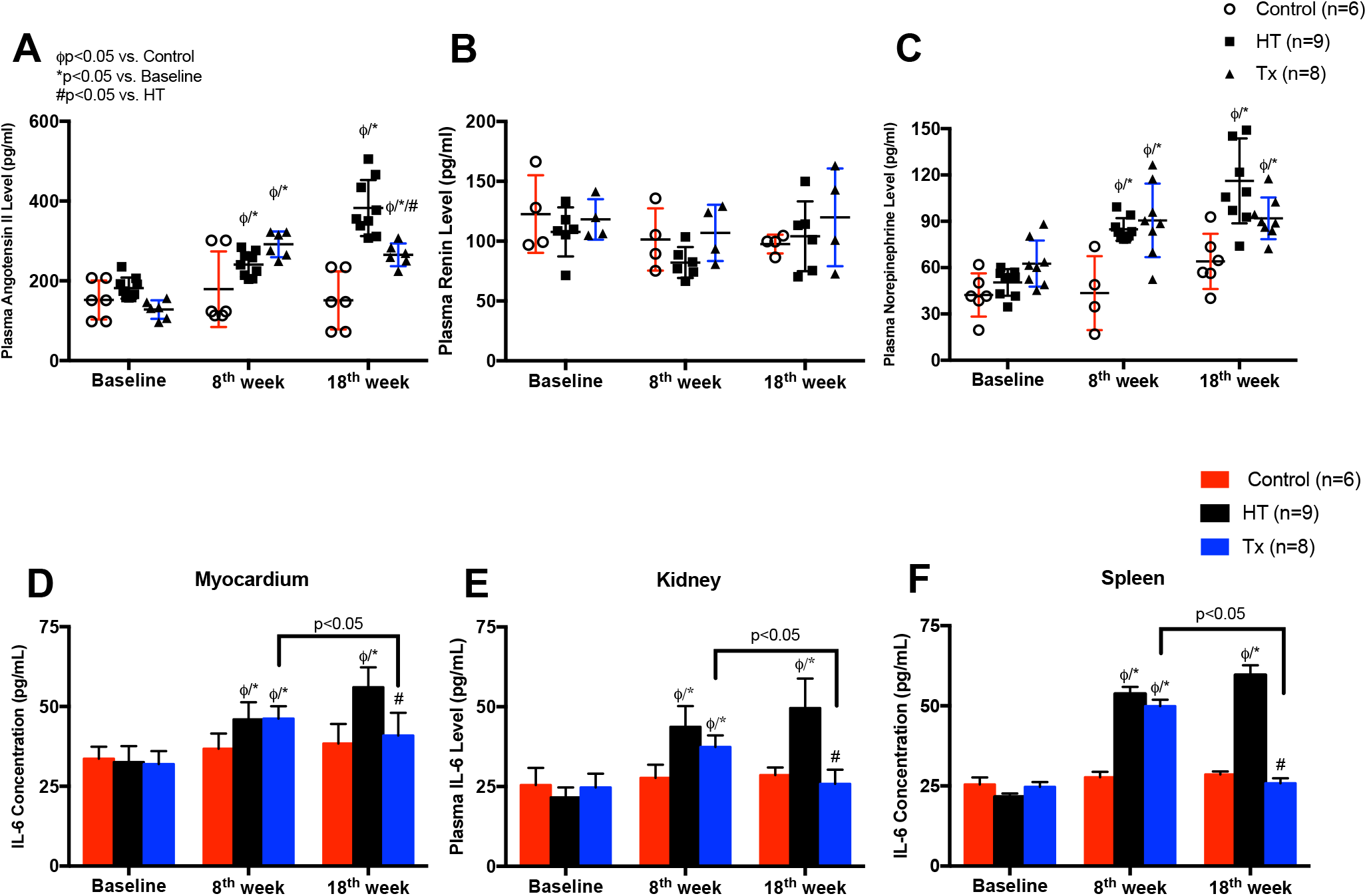
Plasma concentration of biomarkers and tissue concentration of IL-6 in control, hypertension and GLP1RA treatment groups. Serial changes to plasma Ang II (**A**), plasma renin (**B**) and plasma norepinephrine (**C**) as well as IL-6 tissue concentration in myocardium, kidney and spleen (**D-F**) in control (n=6), hypertension (n=9) and GLP1RA treatment (n=8) groups. ϕ<0.05 versus Control; *<0.05 versus Baseline; #<0.05 versus Hypertension.

Exploratory analysis of the changes in the level of pro-inflammatory cytokines after GLP1RA treatment exhibited a significant reduction in IL-6 measured from myocardium, kidney and spleen in Tx group at 18-week follow-up compared with HT group and its 8-week follow-up (**Figure 6D through 6F**; P<0.05). In addition to the data obtained from Masson’s trichrome staining on the fibrosis percentage of myocardial tissue, the concentration of sICAM-1, as a pro-fibrotic and pro-inflammatory molecule, became significantly higher in myocardium, kidney and spleen from pigs in HT group at 8- and 18-week follow-up compared with its baseline, C group and Tx group (**Figure S3D through S3F**; P<0.05). Increased plasma Ang II in our hypertensive pigs ensured an over-activated RAAS initiated by Ang II infusion, accompanied by increased plasma BNP indicating the deteriorated cardiac function. Under such HT condition, systemic SNA became augmented as indicated by increased plasma NE and plasma adrenaline. Over-activated SNS is known to be capable of enhancing inflammation (24), thus local inflammation in our HT animals was significantly severed as expected. Interestingly, GLP1RA treatment in HT-induced SNA elevation and inflammation effectively tuned down RAAS and SNA thereby alleviating the local accumulation of pro-inflammatory cytokines.

## DISCUSSION

In our study, we have successfully established a large animal model with severe HT and HT-induced hCMP using a combination of continuous Ang II infusion and subcutaneous DOCA pellets implantation. After 8 weeks of HT induction, we harvested the pathophysiological features of HT-derived hCMP as evidenced by progressive systolic and diastolic dysfunction on invasive hemodynamic assessment, and significant LV remodeling and cardiac hypertrophy were determined by speckle tracking strain imaging from echocardiography. Our results also illustrated significant local heterogeneity in SNA activation at different organs by measuring the venoarterial NE gradients and tissue content of NE in HT animals. A dramatic increase in NE gradient over myocardium and splanchnic organs after HT induction compared with animals at baseline was observed in our results. However, tissue content of NE only showed significant raise in the spleen. Application of GLP1RA in HT pigs reduced SNA in the spleen, decreased the venoarterial gradient over the splanchnic organs, reduced IL-6 and sICAM-1 levels and attenuated the elevation of BP. In HT pigs with hCMP, treatment of GLP1RA also halted the increase in LVESP and LVEDP as well as the extent of LV remodeling and cardiac hypertrophy. Thus, upon GLP1RA implications both LV systolic and diastolic function were preserved. Although prior studies have successfully induced HT in animals using either a high dose of Ang II infusion (17, 18) or low dose DOCA pellets implantation with a high salt diet (25), severe decrease in peripheral renin activity leading to different degrees of renal injury was observed as an adverse effect of such HT-induction administration. Therefore, in our study we used a combination of low dose continuous Ang II infusion and subcutaneous DOCA pellets to induce sustained and severe elevation of BP. This sustained BP increase in turn was established in a large animal model of HT with hCMP, which is one of the most major causes of HFpEF. Our animal models were featured with gradual rather than acute raise in plasma Ang II without serial changes in plasma renin. Such phenomenon shows high resemblance with the common human phenotype of refractory HT (26). Interestingly, there was no significant change to renal function as measured by plasma creatinine and no evidence of local activation of RAAS over the myocardium or splanchnic organs in our animals.

Even though clinical signs of HF were absent in our hypertensive animals, evidence from echocardiography assessment and speckle tracking strain showed a phenotype of progressive worsening in LV diastolic function and myocardial dysfunction which were pathophysiological features of human hCMP. More characteristics of hCMP like increased LA volume, significant LV hypertrophy and elevated myocardial fibrosis were also detected in our hypertensive animals.

Previous study has investigated the reginal heterogeneity in the activation of SNA during the pathogenesis of the disease in a hypertensive large animal model with high resemblance to human cardiovascular system (21). Our results confirmed the findings in SNA activation of large animal by observing a significant elevation of the venoarterial NE gradient across the myocardium and splanchnic organs as well as tissue content of NE in the spleen after the induction of HT. These findings showed similarity to the observations from clinical (23) and experimental (27) studies in HFrEF, however, our animal models of hCMP with HFpEF are prominent in venoarterial NE gradient across the myocardium.

Sympathetic nerves are believed to play an important role in modulating the immune response (28). Evidence from this study suggests that short-term SNA activation mediates an anti-inflammatory reflect possibly via the regulation of macrophages located in its white pulp to control systemic inflammation. Nevertheless, the long-tern effects of SNA activation on immune response remain elusive. Prior experimental study confirms that the spleen is the key organ involving in linking the nervous system to the immune system (29). This study has speculated that HT activates splenic SNA to prime an immune response that subsequently contributes to the establishment and maintenance of elevated BP. Our results were consistent with postulation as evidenced by a significant increased tissue content of splenic NE in hypertensive animals. In our hypertensive models administrated with GLP1RA treatment, a significant decrease was observed in venoarterial NE gradient across both myocardium and splanchnic organs, and the splenic tissue content of NE. This decrease was associated with a reduced level of pro-inflammatory cytokines IL-6 and pro-inflammatory biomarker sICAM-1. Prior hypertensive large animal study involving splanchnic denervation illustrated similar result by a direct splanchnic denervation surgery which has effectively lowered BP, and NE expression both systemically and locally (21). However, differing from their mechanism in which the reduction in SNA is the initial driving force of BP and inflammation decrease, GLP1RA treatment used in our study may yield the results in an opposite way.

According to the chemical and pharmacological principle of GLP1RA, its function and triggered downstream cellular pathways mimic the protogenetic protein GLP-1 upon GLP-1 receptor activation (30). Previous study using GLP1RA as a potential treatment to either obesity or obesity-derived chronic inflammation showed a significant increase in heart rate of the experimental models (31), and this observation was also detected in our results. Interestingly, GLP1RA possesses the SNA activation property by directly stimulating the sino-atrial node to increase the heart rate (22). Nonetheless, augmented SNA has the potential to exert negative effect on cardiovascular health (32) so that GLP1RA treatment must have improved HT-induced hCMP or HFpEF against its SNA-favoring property in a different way. As a typical treatment of type 2 diabetes, one of the direct functions of GLP1RA is to increase the insulin secretion and decrease the glucagon secretion (33). Changes in these two hormones contribute to a decreased gastric emptying and hepatic production, and eventually they result in body weight loss. Since overweight or obesity is a common cause of chronic inflammation due to the overproduction of pro-inflammatory cytokines such as IL-6 from abnormal or excessive adipose tissue (34), reduced body weight can improve lipid profile, inflammation and decrease the burden on cardiac output. In our results, hypertensive animals receiving GLP1RA treatment showed significant decrease in body weight/body length ratio, implying the weight loss effect of GLP1RA on these models. Accompanied with body weight loss, IL-6 and sICAM-1 as two major cardiovascular-associated pro-inflammatory cytokines exhibited dramatic reduction. Combining our results of both improved inflammation and halted BP increase, HT-induced SNA activation was eventually tuned down. Therefore, as the reduction in SNA is the initial driving force of BP and inflammation decrease in prior study (21), alleviating the HT-derived inflammation might be the key target of GLP1RA treatment in approaching the same BP control and cardioprotective effect on hCMP and HFpEF.

Our study has limitations. We did not observe any significant hemodynamic effect during a specific period after the administration of GLP1RA, so the monitoring of acute effect of this drug is lacking. More noteworthily, GLP1RA is initially used in type 2 diabetes treatment in which insulin secretion and sensitivity is abnormal (35), the safety and efficacy of this drug to patients who are non-diabetic or non-obese but admitting to HT-induced HFpEF alone hence were not determined despite the fact that many patients clinically admitted to type 2 diabetes have acquired cardioprotective effect from GLP1RA (36). Since GLP1RA can directly stimulate sino-atrial node and increase heart rate as an acute effect, we hypothesize that application of this drug to treat HT and HT-induced HFpEF in patients admitted to severe congestive HF should take high considerations.

## PERSPECTIVES

Our study provides novel insight into the therapeutic potential of GLP1RA in HT-induced HF in a large animal model of hCMP. We also unravel the possible underlying mechanism of GLP1RA in BP control and cardio-protection against its SNA activation property.

## SOURCES OF FUNDING

Our study was supported, in part, by Abbott and the Research Grants Council of Hong Kong, General Research Fund.

## DUALITY OF INTEREST

The authors declare no competing conflicts.

## AUTHOR CONTRIBUTIONS

H.S.H. conceived the project and designed the research. Z.Z.Y., L.S.Y., and Z.Z. performed the studies. S.S., L.W.H. and T.A. provided technical support. Z.Z.Y. and H.S.H. analyzed the data and wrote the manuscript.

## Notes

### Competing Interest Statement

The authors have declared no competing interest.

