## Supplementary figures and images for "The effect of glucagon-like peptide-1 receptor agonist (GLP1RA) on hypertensive-induced heart failure with preserved ejection fraction and hypertensive cardiomyopathy"

### Supplementary figures

**Control** **HT** **Tx**

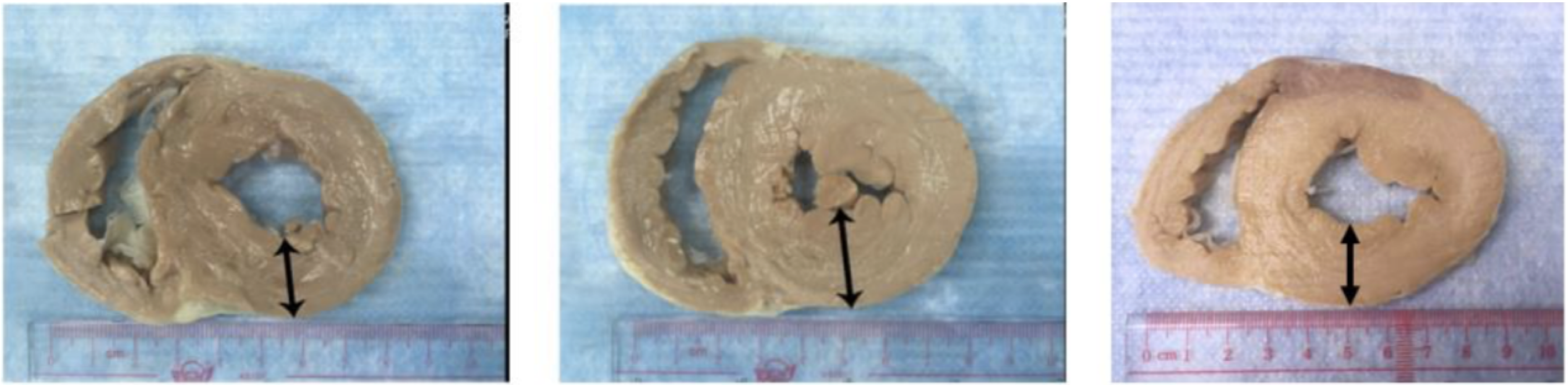


**Supplementary Figure 1**

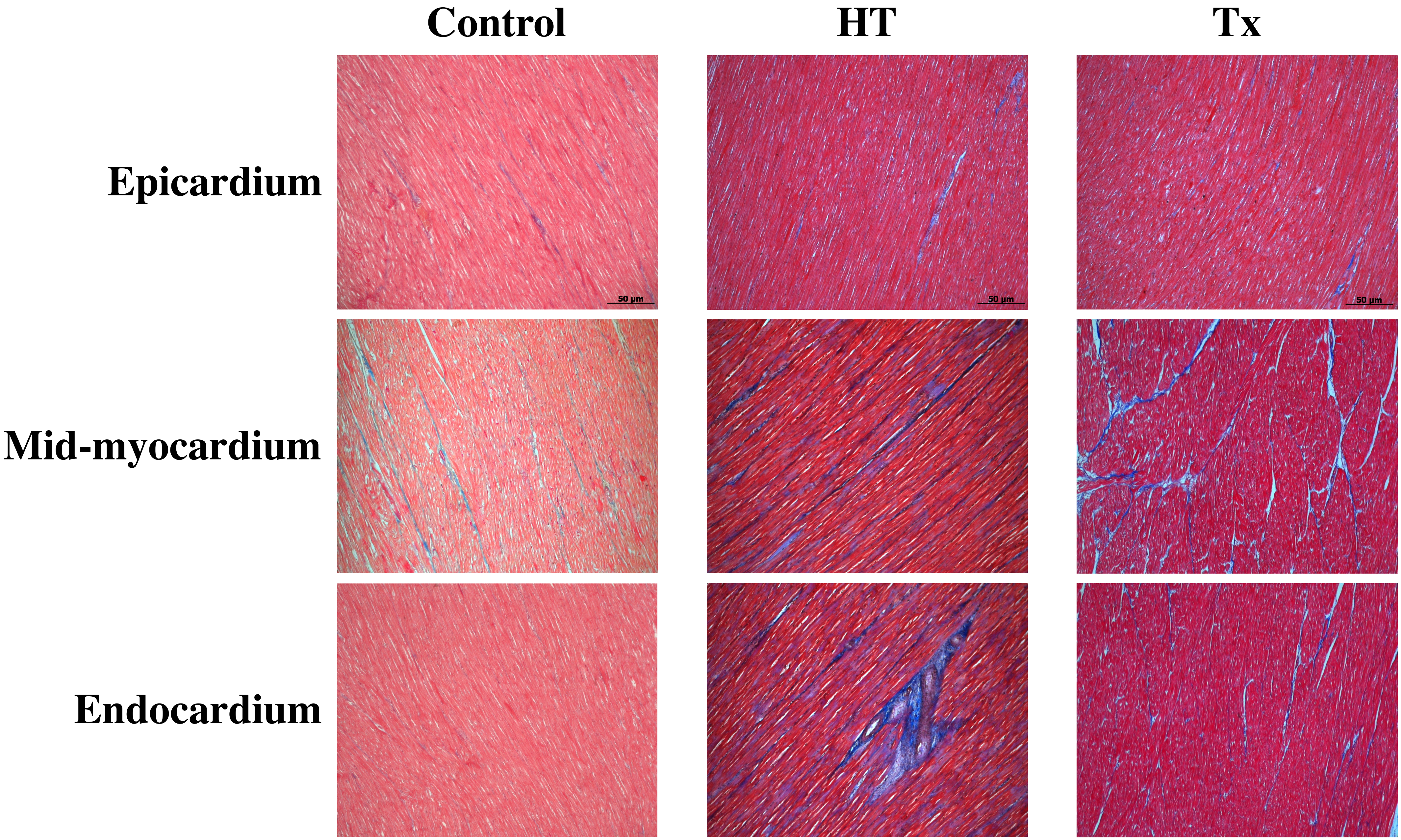


Supplementary Figure 2

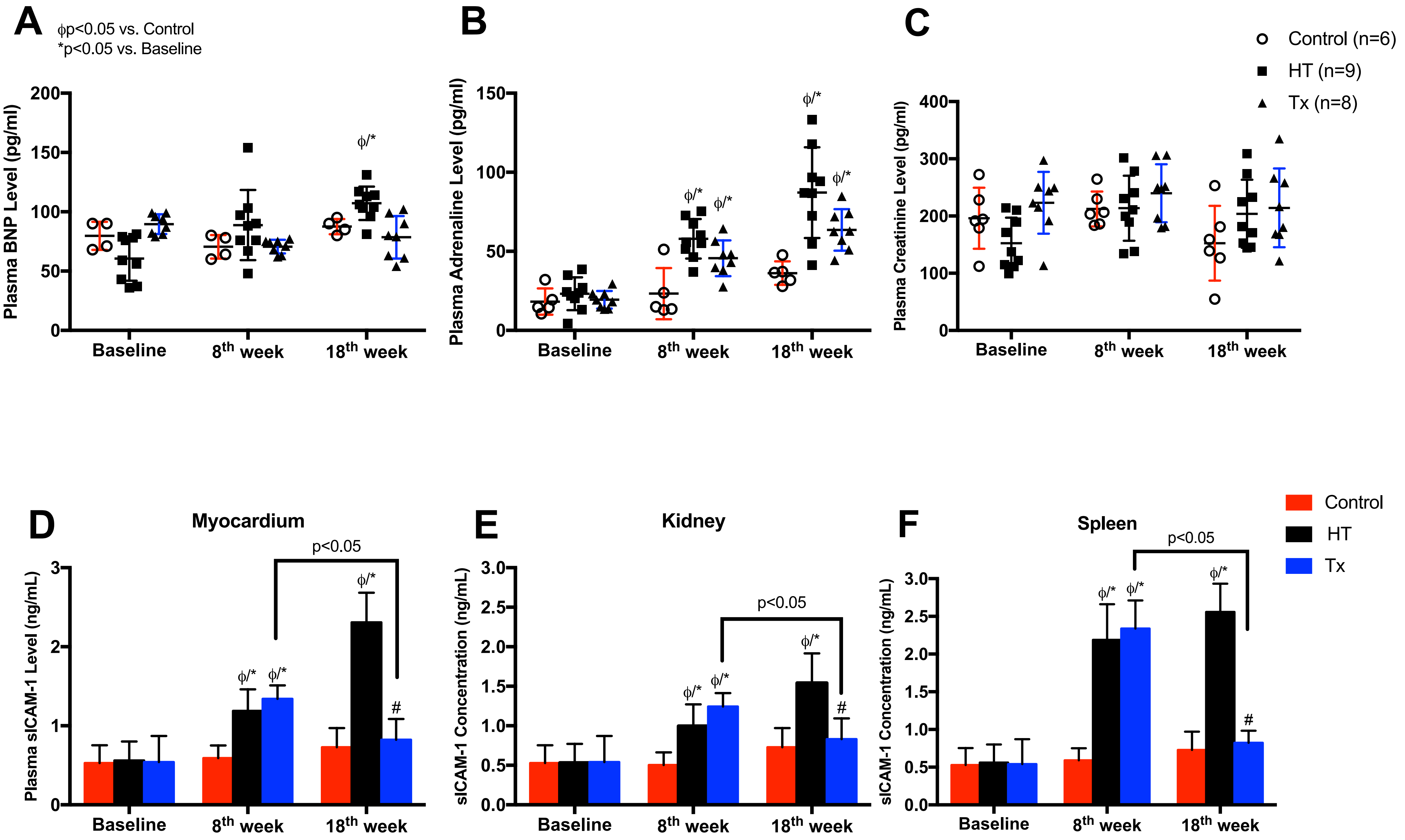


**Supplementary Figure 3**
